# Enhanced Cas12a multi-gene regulation using a CRISPR array separator

**DOI:** 10.1101/2021.01.27.428408

**Authors:** Jens P. Magnusson, Antonio R. Rios, Lingling Wu, Lei S. Qi

## Abstract

The type V-A Cas12a protein can process its CRISPR array, a feature useful for multiplexed gene editing and regulation. However, CRISPR arrays often exhibit unpredictable performance due to interference between multiple crRNAs. Here, we report that Cas12a array performance is hypersensitive to the GC content of crRNA spacers, as high-GC spacers can impair activity of the downstream crRNA. We analyzed naturally occurring CRISPR arrays and observed that repeats always contain an AT-rich fragment that separates crRNAs; we term this fragment a *CRISPR separator.* Inspired by this observation, we designed short, AT-rich synthetic separators (*synSeparators*) that successfully removed the disruptive effects between crRNAs. We demonstrate enhanced simultaneous activation of seven endogenous genes in human cells using an array containing the synSeparator. These results elucidate a previously unknown feature of natural CRISPR arrays and demonstrate how nature-inspired engineering solutions can improve multi-gene control in mammalian cells.

## Introduction

Precise control of cell identity and behavior will require the ability to regulate the expression of many genes simultaneously. Genes can be experimentally turned on or off using transcriptional activators or repressors attached to nuclease-deactivated Cas proteins (dCas) in technologies named CRISPRa and CRISPRi (Lo & Qi, 2017). When this fusion protein is recruited to a target gene promoter or coding sequence, guided by a CRISPR-associated RNA (*crRNA* for type V CRISPR Cas12) or a chimeric single-guide RNA (*sgRNA* for type II CRISPR Cas9), the target gene can be activated or repressed. While this technology works relatively well for controlling single genes, a central goal is to efficiently control more than a handful of genes at a time for applications in cell engineering.

In their native bacterial context, multiple crRNAs are encoded on a CRISPR array transcribed as a single, long transcript (Medina-Aparicio et al., 2017; Pul et al., 2010) (Fig. 1A). Unlike the widely used Cas9, which requires a trans-activating crRNA (tracrRNA) and RNase III for maturation of its guide RNA, Cas12a (also known as Cpf1) can process its own CRISPR array (Deltcheva et al., 2011; Zetsche et al., 2015). Therefore, multiple genes can be controlled by expressing a single CRISPR array together with a Cas12a-based transcription factor. This strategy has recently been used to edit or control the expression of multiple genes in human cells (Breinig et al., 2019; Campa et al., 2019; Kleinstiver et al., 2019; Tak et al., 2017; Zetsche et al., 2017). However, CRISPR arrays encoding multiple crRNAs often exhibit unpredictable performance for multi-gene regulation. Small-RNA-seq experiments suggest that array processing by Cas12a is uneven, such that the constituent crRNAs may differ ten- to hundred-fold in abundance (Campa et al., 2019). To reliably regulate many genes using this system, it will be crucial to optimize the performance of CRISPR arrays in mammalian cells and understand the principles governing CRISPR array processing.

**Figure 1.**
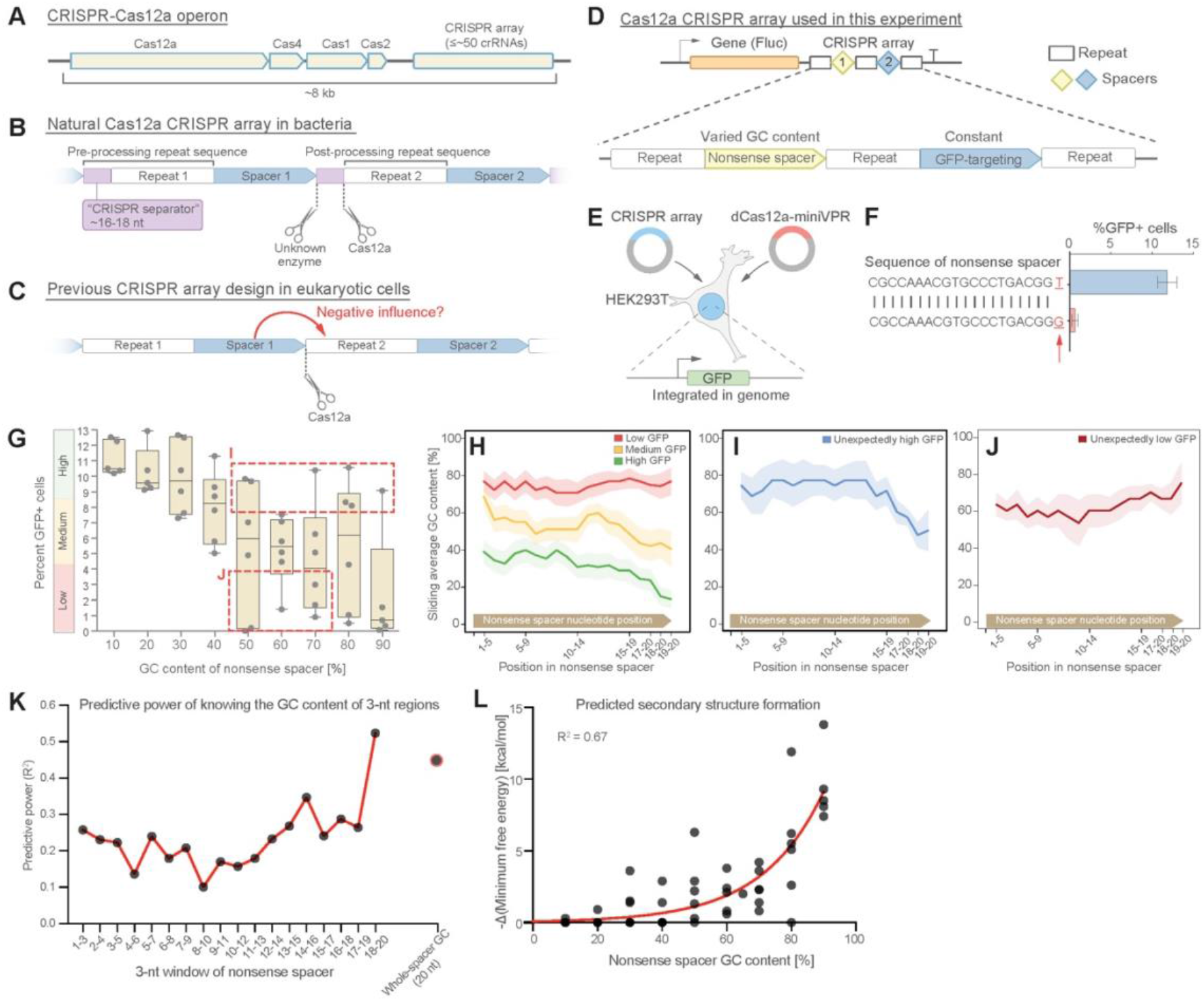
crRNA performance is affected by the GC content of the upstream spacer. (**A**) The CRISPR-Cas12a operon consists of Cas genes and a CRISPR array. (**B**) Each crRNA consists of a repeat and a spacer. Pre-processing repeats contain a ~16-18-nt fragment, here denoted *CRISPR separator,* which gets excised by Cas12a and an unknown enzyme. (**C**) The separator has previously been omitted when expressing Cas12a arrays in mammalian cells. We asked if the separator serves to insulate crRNAs from the negative influence of secondary structure in spacers. (**D**) We designed CRISPR arrays consisting of two crRNAs, the first with a non-targeting nonsense spacer, and the second targeting the promoter of GFP, genomically integrated in HEK293T cells. (**E**) Experimental setup; GFP fluorescence was analyzed as a measure of array performance. (**F**) CRISPR arrays can display hypersensitivity to the composition of the nonsense spacer. In extreme cases, replacing the last nucleotide from T to G can lead to almost complete abrogation of GFP activation. (**G**) A library of 51 CRISPR arrays where the first crRNA contains one nonsense spacer with varying GC content and the second crRNA targeting GFP. A strong negative correlation is seen between the GC content of the nonsense spacer and GFP fluorescence. Each dot represents one of the 51 CRISPR arrays (three replicates). Arrays were divided into three groups based on the level of GFP fluorescence they enabled. Boxes indicate two groups that were analyzed in **I** and **J**. (**H-J**) For each group, the average GC content of a sliding 5-nt window was calculated. The best-performing arrays were the ones where the nonsense spacer happened to have low GC content at its 3’ end. Some arrays showed unexpectedly high or low GFP activity for the GC content of their nonsense spacers (**G**). These arrays contain low (**I**) or high (**J**) GC content at the very 3’ end of their nonsense spacers, suggesting that the GC content of the last few bases is an important predictor of array performance. Shaded regions in **H-J** represent standard error. (**K**) The predictive power of knowing the GC content of 3-nt regions in the nonsense crRNA (Methods). (**L**) A plot showing the relationship between predicted secondary structure (-Δ(minimum free energy) and GC content of the 51 nonsense spacers.

A crRNA consists of a repeat region, which is often identical for all crRNAs in the array, and a spacer, which serves to guide the Cas12a protein to double-stranded DNA with a complementary sequence (Fig. 1B). In their natural bacterial setting, the full-length repeat sequence includes a short (~16-18 nt) fragment that gets excised and discarded during crRNA processing and maturation, leaving the final crRNA to consist of a post-processing repeat and a spacer (Fig. 1B). The excised repeat fragment, which we here denote a *CRISPR separator,* undergoes cleavage at its 3’ end by Cas12a itself, and at its 5’ end by an unknown enzyme (Swarts, 2019) (Fig. 1B). The separator is not strictly needed for array function and has no known role, as far as we are aware. Cas12a cannot excise the separator on its own, which means that it remains attached to the 3’ end of the upstream spacer in mammalian cells (Zetsche et al., 2017). For this reason, the separator has been omitted when Cas12a arrays have been experimentally expressed in eukaryotic cells (Fig. 1C) (Campa et al., 2019; Kleinstiver et al., 2019; Tak et al., 2017; Zetsche et al., 2017). Yet, the presence of a separator sequence in naturally occurring Cas12a CRISPR arrays – despite being under intense selective pressure – suggests that the separator might have a biological function.

Here, we hypothesized that the separator plays a role in facilitating CRISPR array processing and that array performance is impaired in its absence. We find that crRNAs lacking the separator are highly sensitive to the content of guanine and cytosine (GC) nucleotides in the upstream spacer. Specifically, the higher the GC content of a spacer, the worse the downstream crRNA performs. We notice that the natural separator sequence is rich in adenine and thymine/uracil (AT/U) nucleotides, and we therefore surmised that the separator insulates neighboring crRNAs from one another. We design and insert a synthetic separator (*synSeparator)* consisting of 4 A/T nucleotides between each crRNA in a Cas12a CRISPR array, which allows more effective CRISPR activation of multiple endogenous genes in human cells. Based on these results, we conclude that the CRISPR separator found in natural Cas12a arrays acts as an insulator that reduces interference between adjacent crRNAs. These results shed light on the evolutionary forces acting on CRISPR-Cas systems and show how natural systems can guide optimization of CRISPR-mediated multi-gene regulation technology.

## Results

### crRNA performance is affected by the GC content of the upstream spacer

In bacteria harboring CRISPR-Cas systems, new spacers are acquired from viral genomes and integrated into the CRISPR array. It is possible that some spacer sequences form RNA secondary structures that might interfere with Cas12a binding and processing of the array. RNA secondary structure is in fact known to impede Cas protein binding and processing in similar contexts. For example, Cas12a function is sensitive to hairpin formation downstream of the CRISPR array (Liao et al., 2019), and crRNA processing is sensitive to base-pairing of the repeat with regions outside the crRNA (Creutzburg et al., 2020). The RNA-guided, RNA-cleaving protein Cas13 is also negatively affected by crRNA secondary structure (Abudayyeh et al., 2016; Yan et al., 2018). It is therefore theoretically plausible that local secondary structure within the transcribed CRISPR array could interfere with array processing (Fig. 1C). One feature that promotes RNA secondary structure formation is high GC content (Chan et al., 2009), because G-C base pairs contain three hydrogen bonds, compared to A-U’s two hydrogen bonds. We first tested whether the GC content of a spacer could affect performance of a downstream crRNA.

We developed a method for assembling CRISPR arrays using oligonucleotide hybridization and ligation, with which we could assemble CRISPR arrays containing up to 30 crRNAs (Methods). We used this method to assemble a simple Cas12a array consisting of two consecutive crRNAs whose repeat regions did not contain the separator sequence (Fig. 1D). In this array, the second crRNA’s spacer was complementary to the promoter of GFP, which had been genomically integrated into HEK293T cells (Fig. 1E). The first crRNA’s spacer instead consisted of a non-targeting nonsense sequence, which we could vary to study its effect on GFP activation.

To study the consequences of varying the nonsense spacer, we transfected the HEK293T cells with constructs encoding the CRISPR array and nuclease-deactivated Cas12a fused to the miniaturized VP64-p65-Rta activator (Vora et al., 2018) and mCherry (subsequently denoted dCas12a-miniVPR). If the nonsense spacer did not influence the performance of the downstream crRNA, we should see equal GFP activation in all cells. If, on the other hand, the nonsense spacer affected the performance of the downstream spacer, we would measure differences in GFP fluorescence. Surprisingly, we found that the CRISPR array sometimes displayed hypersensitivity to the composition of the nonsense spacer: In extreme cases, a single nucleotide change from T to G led to almost complete abrogation of GFP activation in transfected cells (Fig. 1F, Methods).

We asked how such hypersensitivity could come about. We generated a library of 51 CRISPR arrays containing one nonsense spacer with varying GC content (10-90%), and one GFP-targeting spacer that was identical in all arrays (Table S1). Forty-eight hours after transfection, we analyzed cells by flow cytometry and quantified GFP fluorescence as a measure of array performance. Intriguingly, we observed a strong negative correlation between the GC content of the nonsense spacer and GFP activation (Figs. 1G, S1). This indicated that spacers can exert a strong influence on the performance of the downstream crRNA.

Since these nonsense spacers were random sequences, we asked whether the distribution of GC content within these spacers mattered for their effect on the downstream crRNA. We first divided all the nonsense spacers into three groups based on whether they enabled high, medium or low GFP activation (Fig. 1G). For each nonsense spacer, we calculated the GC content of a sliding 5-nt window (Fig. 1H; Methods). Interestingly, this analysis showed that ”permissive” nonsense spacers had relatively low GC content at the 3’ end, close to the Cas12a cleavage site. In contrast, non-permissive nonsense spacers had slightly higher GC content at the 3’ end. Nonsense spacers that enabled medium-level GFP activation had quite high overall GC content but lower GC content close to the 3’ end. This suggested that the GC content of the spacer’s last few bases, close to the Cas12a cleavage site, might be a more important determinant than the overall GC content of a spacer.

Surprisingly, nonsense spacers in the 50-90% GC range exhibited a wide range of GFP activation, some enabling unexpectedly high GFP activation and others unexpectedly low (Fig. 1G). We analyzed the sliding GC content specifically of these spacers and found that unexpectedly permissive nonsense spacers showed an even stronger trend toward low GC content at the 3’ end (Fig. 1I). In contrast, unexpectedly non-permissive nonsense spacers had high GC content at the 3’ end (Fig. 1J). Thus, even if a nonsense spacer had high GC content, it could still allow efficient performance of the GFP-targeting crRNA if GC content in the last 3-5 bases was low, and vice versa.

GC content of the upstream spacer was moderately predictive of GFP activation (R^2^ = 0.45; Fig. 1K; Methods). Our data suggested that the identity of a spacer’s most 3’ bases might be disproportionately important for the performance of the subsequent crRNA. Indeed, we found that simply knowing the average GC content of the last three nucleotides in the nonsense spacer allowed slightly better predictive power than knowing the GC content of the whole nonsense spacer (R^2^ = 0.52; Fig. 1K). The GC content of these last three bases was more predictive of array performance than that of any other three bases in the nonsense spacer (Fig. 1K). These results indicated that high GC content at the 3’ end of a spacer impairs performance of the subsequent crRNA.

GC content is a determinant of secondary structure formation. We used the online tool RNAfold (Lorenz et al., 2011) to calculate secondary structure formation for the library of 51 nonsense spacers. We indeed found a strong positive correlation between GC content and predicted secondary structure formation (R^2^ = 0.67; Fig. 1L). Taken together, these results suggest that high GC content in spacers can lead to secondary structure formation, which may interfere with proper crRNA processing by Cas12a.

### Natural CRISPR arrays contain separator sequences with low GC content

We next asked whether bacteria have evolved mechanisms to preferentially incorporate low-GC spacers into their CRISPR arrays to overcome the hypersensitivity to spacer GC content. After all, it could be detrimental to a bacterium if it accidentally incorporated a high-GC spacer that lowered the performance of a pre-existing crRNA. To address this question, we analyzed 727 naturally occurring Cas12a spacer sequences from 30 bacterial species containing the Type V-A CRISPR (Table S2; Methods). However, we did not find any conspicuous absence of GC-rich spacers: Spacer GC content was distributed around an average of 39%, with a range of 10-70% (Fig. 2A), though spacer GC content did vary between species and was weakly correlated with the overall genomic GC content (R^2^ = 0.16; Fig. S2A). These results were true also for *Lachnospiraceae bacterium,* the species from which our Cas12a variant was derived, and for the commonly used *Acidaminococcus sp.* and *Francisella tularensis* subsp*. novicida* (Fig. 2A). Neither did we find that GC content was lower at the 3’ end of these spacers (Fig. 2B; Methods). We therefore wondered how natural CRISPR arrays cope with high-GC spacers.

**Figure 2.**
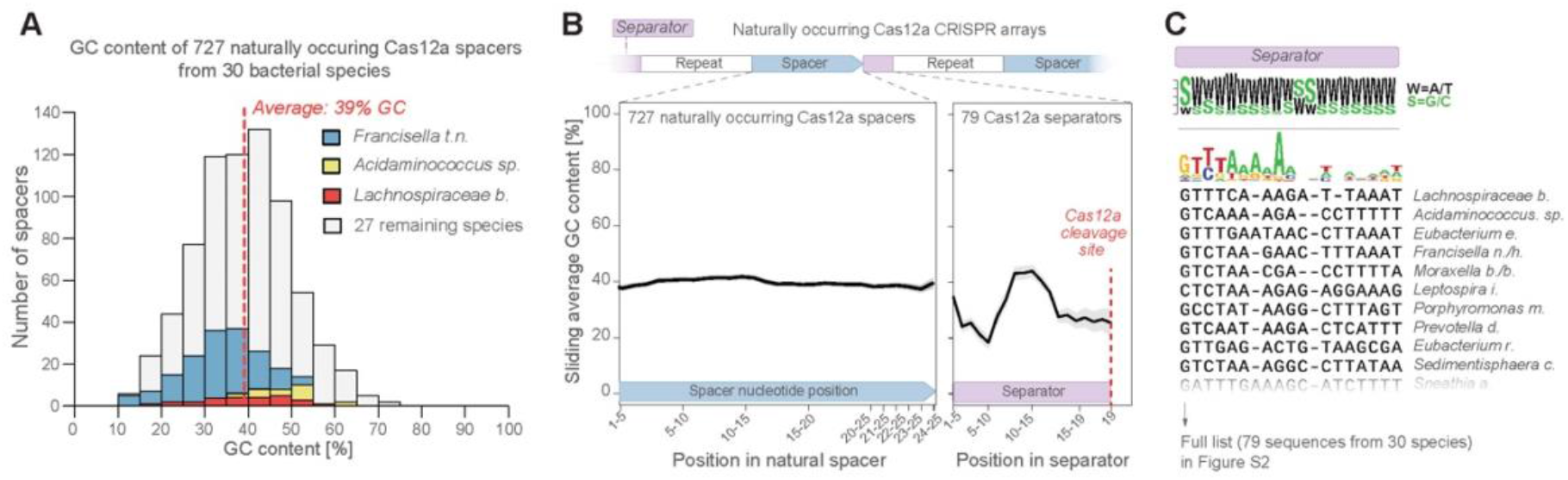
Natural CRISPR arrays contain separator sequences with low GC content. (**A**) The GC content of naturally occurring CRISPR-Cas12a spacers display no obvious depletion of high-GC spacers (though spacer GC content is weakly correlated with overall genomic GC content [Fig. S2A]). Spacers from commonly used Cas12a variants (Lb, As, Fn) are highlighted in color. (**B**) Neither do these spacers show low GC content at their 3’ ends. But the separator sequences of these crRNAs have low GC content. (**C**) This is seen also in a multiple-sequence alignment of 79 natural separator sequences, which suggests that one purpose of the CRISPR separator is to act as an insulator between adjacent crRNAs in a Cas12a CRISPR array.

In naturally occurring CRISPR arrays, the separator sequence gets excised through the action of Cas12a and an unknown enzyme (Fig. 1B) (Zetsche et al., 2015). We asked whether the separator might act as an insulator that protects every crRNA from disturbances caused by the secondary structure in upstream spacers. We analyzed 79 unique separator sequences from 30 bacterial species (Table S2; Methods). Overall, these separators showed little sequence conservation (Figs. 2C, S2B), except for a moderately conserved region at the very 5’ end (GTYTA). This conserved region possibly acts as a recognition motif for the unknown enzyme responsible for cleaving the separator at the 5’ end. However, we did detect a strong bias for low GC content in the 79 separator sequences (Fig. 2B-C). This opened the possibility that in natural CRISPR arrays, the separator aids CRISPR array processing by providing an AT-rich sequence that maximizes Cas12a accessibility to its cleavage site.

Like Cas12a, the type VI CRISPR RNA-cleaving enzyme Cas13d can also process its own crRNA (Konermann et al., 2018; Yan et al., 2018; B. Zhang et al., 2019; C. Zhang et al., 2018), an ability that makes Cas13d attractive for multi-gene regulation on the RNA level (Fig. S2C). We similarly analyzed the 6-nt CRISPR separator from 12 bacterial species containing Cas13d arrays (Table S2; Methods). We found a strong bias for low GC content here as well, particularly at the 3’ end of the separator, close to the Cas13d cleavage site (Fig. S2C). This suggests that sensitivity to spacer GC content is a factor that has shaped the evolution of multiple CRISPR-Cas systems.

### A synthetic separator improves CRISPR array performance in human cells

We asked whether Cas12a CRISPR arrays would show improved performance in human cells if they included the natural CRISPR separator between each crRNA. To test this, we designed CRISPR arrays where the first crRNA contained a nonsense spacer (30% GC content) and the second a GFP-targeting spacer, with and without the natural, 16-nt separator from *L. bacterium* preceding each repeat (Fig. 3A; Methods). Including this separator, however, almost completely abolished the array’s function, as almost no GFP activation was seen in the transfected cells (Fig. 3B). This is consistent with a previous study that reported poor performance of crRNAs containing the pre-processed Cas12a repeat, which contains the separator (Liu et al., 2019), and is why previous designs have completely excluded any separator sequence. One possible reason is that the long separator remains attached at the 3’ end of each spacer because Cas12a cannot fully excise it (Zetsche et al., 2015, 2017), which might interfere with Cas12a function, as has been shown previously (Nguyen et al., 2020).

**Figure 3.**
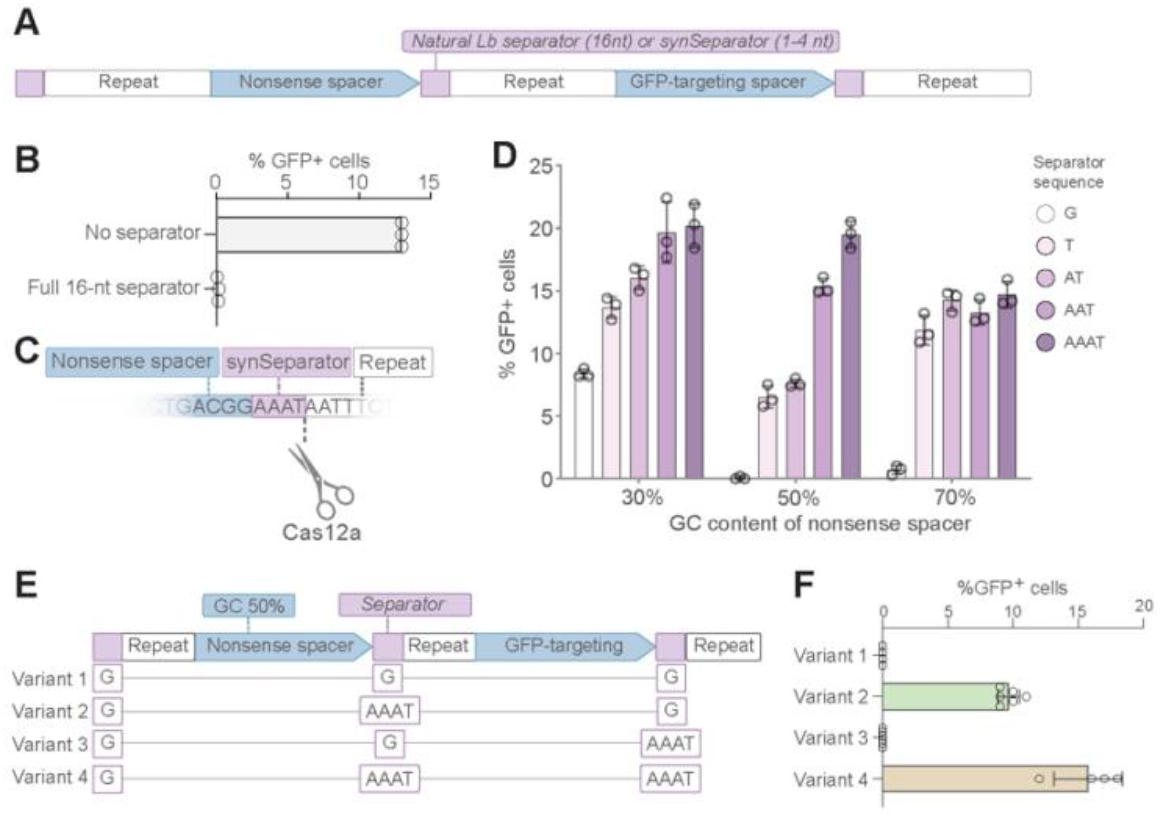
A synthetic separator improves CRISPR array performance in human cells. **(A)** A diagram showing how we introduced either the full, 16-nt separator sequence from *L. bacterium* or a short 1-4 nt synthetic separator (synSeparator) between each crRNA in the CRISPR array. (**B**) The full, 16-nt separator sequence almost completely eliminates GFP activation. (**C**) Diagram showing inclusion of synSeparator sequences from the 3’ end of the *L. bacterium* in the CRISPR array. Each array contained a synthetic separator (G, T, AT, AAT, or AAAT) upstream of every repeat. The GC content of the nonsense spacer was 30%, 50% or 70%. The most non-permissive 50- and 70%-GC spacers from Fig. 1G were used. (**D**) AT-rich synSeparators improve GFP activation compared with a single G nucleotide. (**E-F**) This effect is caused by the separator upstream of the GFP-targeting crRNA: Adding the synSeparator only downstream of the GFP-targeting crRNA leads to no improvement. This suggests that the separator acts by improving crRNA processing rather than by generating an AAAT overhang on the GFP-targeting spacer itself.

We instead asked if we could incorporate only a portion of the separator and still retain its insulating function. We generated CRISPR arrays in which all crRNAs were either preceded by 1-4 A/T nucleotides from the natural *L. bacterium* separator, or by a single G as a control (Fig. 3A, C). We generated three versions of each array, where the GC content of the upstream spacer was 30%, 50% or 70%. (We used the 50% and 70% spacers that had resulted in the lowest level of GFP activation in Fig. 1G) (Table S3). Interestingly, addition of an AT-rich synthetic separator improved the performance of the CRISPR array in all cases (Fig. 3D), suggesting that a very short, AT-rich synthetic separator was sufficient to counteract the interference between crRNAs. We denoted this new sequence a *synSeparator* for synthetic separator.

CrRNA performance can be increased by adding RNA extensions to the 3’ end of the crRNA’s spacer (Creutzburg et al., 2020; Kocak et al., 2019; Nguyen et al., 2020). We asked whether the increased GFP activation we observed with the AAAT synSeparator was caused by the AAAT extension added to the 3’ end of the GFP-targeting spacer, rather than by the AAAT separating the two crRNAs in this array. We made four variants of this array where we changed the position of the G or AAAT separators (Fig. 3E). We observed activated GFP expression with *variant 2* but not with *variant 3* (Fig. 3F), supporting the hypothesis that the synSeparator exerts its main effect by facilitating crRNA insulation and processing. We did observe a small additional increase in effectiveness using *variant 4* compared to *variant 2.* This can either be attributed to improved insulation and processing at the 3’ end of the GFP spacer itself (which has 45% GC content), or to increased performance of the GFP-targeting spacer due to the 3’ AAAT extension on this spacer.

### The synthetic separator improves multiplexed gene activation in human cells

We next tested if the addition of the synSeparator would improve CRISPR activation of endogenous genes when crRNAs were expressed in a CRISPR array. We transfected HEK293T cells (Fig. 1E) with a CRISPR array containing seven crRNAs (Fig. 4A), each targeting the promoter of a different gene, and a dCas12a-VPR-mCherry activator (Chavez et al., 2015). Target genes included both protein-coding genes (*CD9, IFNG, EGFR, TMEM107, GFP)* and long non-coding RNA (lncRNA) genes (*IMPACT, DANCR)* (Table S3; Methods). The seven target genes differed in their baseline expression levels in HEK293T cells (Fig. 4B). We analyzed target gene levels using RT-qPCR on bulk-RNA isolates and did not sort transfected cells based on uptake of the Cas12a or CRISPR array plasmids.

**Figure 4.**
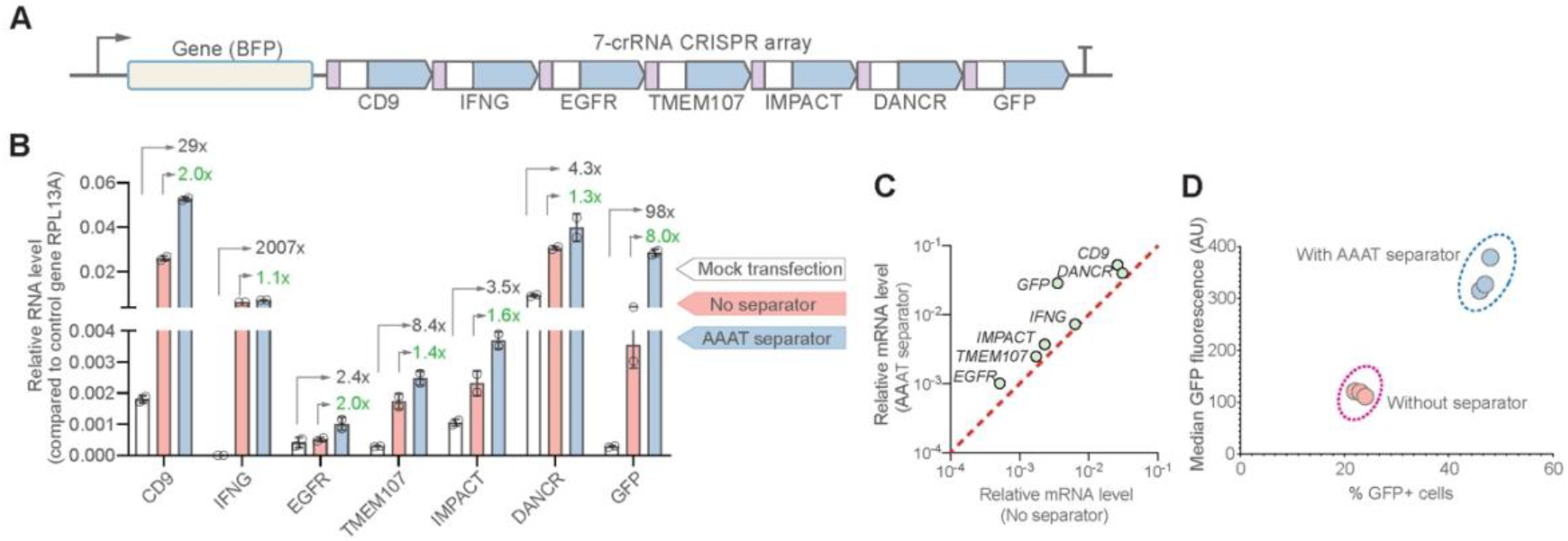
The synthetic separator improves multiplexed gene activation in human cells. (**A**) We designed a 7-crRNA array to activate seven endogenous/reporter genes in HEK293T cells with and without the AAAT synSeparator between each crRNA. (**B**) For all target genes, the synSeparator improves target gene activation level, as measured by RT-qPCR. Green values show the fold-change difference with and without the synSeparator. (**C**) A plot showing that the improvement of CRISPR activation is consistent across genes. (**D**) This improvement is also seen on the protein level for the target gene GFP, as measured by GFP fluorescence and percent GFP-positive cells.

Including the synSeparator significantly increased activation levels of all target genes compared to the array without the synSeparator (Fig. 4B). The improvement effect varied, ranging from 1.1-fold to 8.0-fold, but was consistent across all target genes (Fig. 4C). The increase was also seen on the protein level, which we could analyze for GFP (Fig. 4D). These results indicated that addition of a short, AT-rich sequence between crRNAs in a CRISPR array consistently improved CRISPR activation performance in mammalian cells.

## Discussion

In this study, we found that Cas12a CRISPR arrays are sensitive to spacer GC content. Specifically, high GC content at the 3’ end of a spacer leads to poor performance of the subsequent crRNA. This effect can be counteracted if the crRNAs are separated by short, AT-rich sequences. Naturally occurring crRNAs do contain such AT-rich sequences, which we here denote *CRISPR separators.* While the full-length, natural separator renders CRISPR arrays non-functional in mammalian cells, we found that a short, synthetic separator can insulate crRNAs from the negative influence of upstream spacers. The simple addition of such a synSeparator reliably improves performance of Cas12a CRISPR arrays for multiplexed gene regulation.

We speculate that the synSeparator acts by reducing secondary-structure formation close to the Cas12a cleavage site and thereby facilitates Cas12a binding and crRNA processing. Spacer GC content is correlated with predicted secondary structure formation (Fig. 1L), and GC content close to the Cas12a cleavage site is particularly predictive of the performance of the downstream crRNA (Fig. 1K). Furthermore, previous data show that Cas12a processing is sensitive to RNA secondary structure (Creutzburg et al., 2020).

One alternative explanation for why the synSeparator improves CRISPR array performance is that the synSeparator improves the GFP-targeting spacer itself rather than facilitating crRNA processing. In other words, that it is the synSeparator downstream of the GFP-targeting spacer, rather than the one upstream, that improves performance by remaining attached to the post-processed GFP-targeting spacer. This would be in line with studies showing that crRNA performance can be improved by RNA 3’ spacer extensions (Creutzburg et al., 2020; Kocak et al., 2019; Nguyen et al., 2020). However, our results demonstrate that the synSeparator exerts its beneficial effect before or during crRNA processing, rather than by modifying the 3’ end of the post-processed GFP-targeting spacer itself (Fig. 3F; *version 2* vs. *version 3*). Although we cannot exclude that a 3’ extension to the GFP-targeting crRNA itself contributes to crRNA efficacy (Fig. 3F, *version 3* vs. *version 4),* such an effect would be marginal.

Yet another alternative explanation for the synSeparator’s function could be that the natural and synthetic separators provide a specific sequence motif that aids Cas12a binding and/or processing rather than counteracting secondary structure formation. However, natural separators contain no conserved sequence close to Cas12a’s binding site that could act as such a binding motif (Fig. 2C), making this possibility unlikely.

We find evidence that high-GC spacers are more likely to form secondary structures (Fig. 1L), but it is theoretically possible that spacer GC content influences array performance through other mechanisms. For example, could some spacers promote RNA polymerase detachment and incomplete transcription of the array gene? We believe that this is unlikely: CRISPR array performance was not correlated with an inadvertent introduction of polyadenylation sites (Beaudoing et al., 2000) in the nonsense spacer, which would serve to prematurely terminate the CRISPR array transcript. In any case, polyadenylation sites are AT-rich, so if this was an issue, AT-rich nonsense spacers would be expected to perform worse than GC-rich ones, which is the opposite of what was observed (Fig. 1G).

Applications involving CRISPRa would benefit from an ability to control target gene expression levels. However, this is difficult with current technologies. The synSeparator could potentially be used as a regulator for tuning target gene activation levels, as crRNA performance can be modulated by modifying the length and nucleotide composition of the synSeparator (Fig. 3D).

It is informative to compare the separator of Cas12a with that of Cas13d. Cas13d’s pre-processed repeat also contains an AT-rich separator sequence. Like Cas12a, Cas13d itself can only cleave the 3’ end of its separator (Yan et al., 2018). But Cas13d crRNAs nonetheless perform well in mammalian cells despite the separator remaining attached to the spacer. Perhaps this is because the Cas13d separator is only 6 nt long, in contrast to Cas12a’s 16-18 nt. A recent study found that the performance of Cas12a crRNAs starts to drop if RNA sequences >13 nt long are attached to the 3’ end of the spacer (Nguyen et al., 2020).

The bacterial enzyme responsible for processing the 5’ site of the natural separator is not known. Cas12a crRNAs do get processed properly when Cas12 and its CRISPR array are ectopically expressed in *E. coli* (Zetsche et al., 2015). We think that the enzyme is unlikely to be a Cas protein because there is no single Cas protein that is always present together with Cas12a in the Cas operon: In the 30 bacterial Cas operons we analyzed, Cas1 was present in 78%, Cas2 in 84% and Cas4 in 78%. The responsible enzyme may be one of the ≥16 RNases present in *E. coli* that performs a similar function in its native bacterium host.

As a biotechnology, improvement of simultaneous gene targeting is beneficial to several applications. In this work, we demonstrated improvement of endogenous gene activation using the redesigned CRISPR array containing the synthetic separator, but it is possible that the same design can be used for multiplexed gene editing and base editing using the Cas12a system. Furthermore, because CRISPR-mediated multi-gene activation has been increasingly used for stem cell reprogramming and large-scale genetic screening (Replogle et al., 2020; Weltner et al., 2018), our method will further enhance these emerging applications. Taken together, the results from our study suggest that the CRISPR-separator has evolved to act as an insulating mechanism that protects crRNAs from the disrupting effects of varying GC content in upstream spacers. Furthermore, our study demonstrates how design inspired by the natural CRISPR system can improve the efficacy of tools for CRISPR gene editing and regulation.

## Acknowledgments

We thank Hannah Kempton for vectors encoding Cas12a protein. J.P.M. is supported by the Human Frontier Science Program Long-term Fellowship and the Sweden-America Foundation. S.L.Q. is supported by Li Ka Shing Foundation and NIH the National Institutes of Health Common Fund 4D Nucleome Program (U01 EB021240).

## Author Contributions

J.P.M. designed the study, performed experiments, analyzed results and prepared the manuscript. A.R.R. and L.W. performed experiments. S.L.Q. performed project design, coordination and supervision, interpreted data and prepared the manuscript.

## Competing interests

The authors have filed a provisional patent application related to this work.

## Materials and Methods

### Cell culture

We used HEK293T cells (Clontech, Mountain View, CA) carrying a genomically integrated dscGFP gene driven by the TRE3G promoter (consisting of seven repeats of the Tet response element) (Kempton et al., 2020). This cell line was clonally sorted and expanded and showed no background GFP fluorescence. Cells were cultured in DMEM + GlutaMAX (Thermo Fisher, Waltham, MA) containing 100 U/mL of penicillin and streptomycin (Thermo Fisher) and 10% Fetal Bovine Serum (Clontech). Cells were grown at 37°C with 5% CO_2_ and passaged using 0.05% Trypsin-EDTA solution (Thermo Fisher) or TryplE Express Enzyme (Thermo Fisher).

### Transfection

Cells were transfected with constructs carrying 1) the nuclease-deactivated (D832A) dCas12a (from *Lachnospiraceae bacterium,* human codon-optimized) (Zetsche et al., 2015) fused either to the VP64-p65-Rta (VPR) activator (Chavez et al., 2015) and mCherry, or to mini-VPR (Vora et al., 2018) and mCherry; 2) a CRISPR array-expressing plasmid. For Figures 1 and 3, a CRISPR array construct consisting of firefly luciferase immediately followed by a CRISPR array and an SV40 pA terminator, expressed under the CAG promoter element, was used (Supplementary File 1). For the activation of seven endogenous genes (Fig. 4), firefly luciferase was replaced with BFP and a Malat1 Triplex sequence (Campa et al., 2019) followed by the *L. bacterium* Cas12a leader sequence (Supplementary File 1).

Cells were seeded one day before transfection at a density of 5×10^4^ cells per well in a 24-well plate. Cells were transfected using TransIT-LT1 transfection reagent (Mirus Bio, Madison, WI) according to the manufacturer’s recommendation (250 ng dCas12a-VPR-mCherry plasmid; 250 ng CRISPR array plasmid; 1.5 μl transfection reagent per well).

### Flow cytometry

Two days after transfection, cells were dissociated using 0.05% Trypsin-EDTA or TrypLE (Thermo Fisher), passed through a 40 μm filter-capped test tube (Corning, Corning, NY), and analyzed using either a CytoFLEX S flow cytometer (Beckman Coulter, Brea, CA) or a BD Influx FACS machine (BD Biosciences, Franklin Lakes, NJ) or a BD FACSMelody (BD Biosciences). For each experiment, 10,000 events were recorded. During flow cytometry analysis, we gated for cells expressing the Cas12a construct (mCherry^+^) but not the CRISPR array construct. That is because array processing by Cas12a severs the upstream reporter gene from the poly-A tail, thus potentially disturbing reporter gene expression and thereby the analysis. For Figs. 1 and 3, three replicates were performed for each sample.

### RT-qPCR to quantify endogenous gene activation (Fig. 4)

Cells were transfected as described above. For cell harvesting, all cells in each well were dissociated and included in the analysis and were thus not sorted based on uptake of Cas12a or CRISPR array plasmids. Two biological replicates were performed. Total RNA was extracted with the RNeasy Plus Mini Kit (Qiagen, Germany), according to manufacturer’s instructions. Reverse transcription was performed using iScript cDNA Synthesis kit (Bio-Rad, Hercules, CA). Quantitative PCR reactions were run on a LightCycler thermal cycler (Bio-Rad) with iTaq Universal SYBR Green Supermix (Bio-Rad). ΔΔCt values for the target genes were divided by those of *RPL13A* to obtain relative expression. crRNA spacers and RT-qPCR primers are listed in Table S3.

### Assembly of CRISPR arrays

CRISPR arrays were assembled using an oligonucleotide duplexing and ligation method that we developed. First, arrays were designed computationally using SnapGene software (v. 5.1-5.2; Insightful Science, San Diego, CA). The arrays were designed to include two flanking sequences containing a 20-bp overlap with the opened backbone plasmid, as required for a subsequent In-Fusion reaction. This double-stranded CRISPR array sequence was then computationally divided into ≤60-nt DNA sequences with unique 4-nt 5’ overhangs, which were ordered from Integrated DNA Technologies (IDT, Coralville, IA) in LabReady formulation (100 μM in IDTE buffer, pH 8.0) and standard desalting purification. For each ligation vial, an *oligonucleotide mix* was first made containing 1 μl of each oligonucleotide. Up to 16 single-stranded oligonucleotides (i.e. corresponding to 8 oligo duplexes) were ligated per reaction vial. For CRISPR arrays longer than that, the reaction was divided into multiple vials, each vial containing ≤8 oligonucleotide duplexes (e.g. if the array consists of 12 oligonucleotide duplexes, the reaction was performed in two vials with 6 duplexes in each).:

#### Phosphorylation and duplexing

|  |  |
| --- | --- |
| Oligonucleotide mix | 1.0 μl |
| 2x T7 ligation buffer (NEB) | 2.5 μl |
| H<sub>2</sub>O | 1.25 μl |
| <u>T4 PNK (NEB)</u> | <u>0.25 μl</u> |
| *Total* | *5 μl* |

#### Then run a phosphorylation-duplexing reaction on a thermocycler

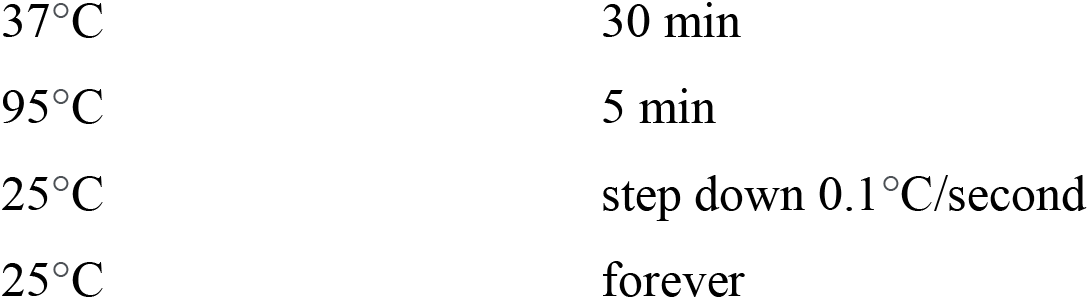

Then, add 1 reaction volume (5 μl) of 1x T7 buffer (2.5 μl 2x T7 buffer + 2.5 μl water). Add 1 μl T7 DNA ligase (New England Biolabs, MA, USA) (Important: Use T7 ligase rather than T4 ligase, as T7 ligase lacks the ability to ligate blunt ends). Incubate at 25°C for 3 hours. Then, dilute the sample 1/5 by adding 40 μl water. Run the sample on a 2% agarose gel. A ladder pattern should be visible. Excise the band corresponding to the ligated product. Depending on whether the entire CRISPR array was assembled in a single vial, or divided into several vials, do either of the following:

#### If the entire array was assembled in a single vial

Gel-purify the excised band using the Macherey-Nagel NucleoSpin Gel & PCR Clean-up kit (Macherey-Nagel, Germany). Insert the purified array into the opened plasmid backbone using In-Fusion cloning (Takara Bio, Japan).

#### If the array was divided into >1 vial

For all excised bands belonging to the same array, pool the excised bands into a single vial. Gel-purify the pooled bands using the Macherey-Nagel NucleoSpin Gel & PCR cleanup kit. Elute in 15 μl water. Then, add 1 volume (15 μl) of 2x T7 buffer and 1 μl T7 DNA ligase. Incubate at 25°C for 3 hours. Then, run the ligated product on a 2% agarose gel. A faint band should be seen corresponding to the full-length CRISPR array. Excise and gel-purify this band. Insert into backbone vector using In-Fusion.

Design of short CRISPR arrays (2 crRNAs) for testing effect of GC content of upstream spacer (Fig. 1)

The 51 nonsense spacer sequences (Fig. 1) were adapted from a negative-control sgRNA library generated by Gilbert et al. (Gilbert et al., 2014). These sequences correspond to scrambled Cas9 spacer sequences, and we adjusted them slightly for length (20 nt) and GC content. All nonsense spacers are listed in Table S1.

#### Computation of GC content in sliding window (Figs. 1H-J, 2B)

For each of the nonsense spacer sequences, we computed the GC content in a sliding 5-nt window (first nucleotides 1-5, then nucleotides 2-6, etc.). For each such window, we then calculated the average and standard error of all 51 spacers. As the sliding window approached the 3’ end of the spacers, we reduced the size of the sliding window to 4, then 3, then 2 nucleotides, in order to increase resolution at the very 3’ end. This was performed also for naturally occurring spacers (Fig. 2B) and CRISPR separators (Fig. 2B). The spacers we analyzed varied in length from 25-36 nt. For this analysis, we truncated the 5’ ends of spacers longer than 25 nt. This way, we could align and analyze the 25 nucleotides at the most 3’ end of every spacer, even though it meant that we would lose information at the 5’ end of longer spacers. For the separator sequences, we first aligned them using the T-Coffee alignment tool (see below), which did not truncate any of the separator sequences.

#### Calculation of the predictive power of spacer GC content

For calculating the predictive power of knowing the GC content of 3 bases in the nonsense spacer (Fig. 1K), we divided each 20-nt spacer into 18 3-nt windows and calculated the GC content for each window. For each such window (e.g. nucleotides 1-3), we plotted GC content versus *percent GFP^+^ cells* for all 51 arrays. We then performed a linear regression (GraphPad Prism v. 9.0; GraphPad Software, San Diego, CA) and used the resulting R^2^ value for Fig. 1K.

#### Multiple sequence alignment of naturally occurring CRISPR sequences (Figs. 2, S2)

To find bacterial CRISPR-Cas12a operons, we used CRISPR-Cas++ (Couvin et al., 2018) using two search strategies: 1) Using the CRISPRCasdb-Blast tool with default settings (accessed September 2020), we input the Cas12a repeat sequences from Zetsche et al. (Zetsche et al., 2015) and extracted all spacers and repeats, making sure that the spacer sequences were all directed in the 5’-to-3’ direction. 2) Using the CRISPRCasdb function, we searched for Cas12a loci in all organisms using default settings. Alignment of separator sequences and post-processed repeats was performed using the multiple sequence alignment tools of SnapGene (v. 5.1-5.2). The separator sequences were aligned using the T-Coffee algorithm. The sequences used can be found in Table S2.

