## Supplemental File 1 for "Enhanced Cas12a multi-gene regulation using a CRISPR array separator"

CAGp-FireflyLuciferase-Array-Terminator (Fig. 3C)

CAG promoter

Firefly luciferase

CRISPR array (AAAT separator, Repeat, 50%-GC nonsense spacer and GFP-targeting spacer)

SV40 terminator

GTGAGCCCCACGTTCTGCTTCACTCTCCCCATCTCCCCCCCCTCCCCACCCCCAATTTTGTATTTATTTATTTTTTAATTATTTTGTGCAGCGATGGGGGCGGGGGGGGGGGGGGCGCGCGCCAGGCGGGGCGGGGCGGGGCGAGGGGCGGGGCGGGGCGAGGCGGAGAGGTGCGGCGGCAGCCAATCAGAGCGGCGCGCTCCGAAAGTTTCCTTTTATGGCGAGGCGGCGGCGGCGGCGGCCCTATAAAAAGCGAAGCGCGCGGCGGGCGGGAGTCGCTGCGTTGCCTTCGCCCCGTGCCCCGCTCCGCGCCGCCTCGCGCCGCCCGCCCCGGCTCTGACTGACCGCGTTACTCCCACAGGTGAGCGGGCGGGACGGCCCTTCTCCTCCGGGCTGTAATTAGCGCTTGGTTTAATGACGGCTCGTTTCTTTTCTGTGGCTGCGTGAAAGCCTTAAAGGGCTCCGGGAGGGCCCTTTGTGCGGGGGGGAGCGGCTCGGGGGGTGCGTGCGTGTGTGTGTGCGTGGGGAGCGCCGCGTGCGGCCCGCGCTGCCCGGCGGCTGTGAGCGCTGCGGGCGCGGCGCGGGGCTTTGTGCGCTCCGCGTGTGCGCGAGGGGAGCGCGGCCGGGGGCGGTGCCCCGCGGTGCGGGGGGGCTGCGAGGGGAACAAAGGCTGCGTGCGGGGTGTGTGCGTGGGGGGGTGAGCAGGGGGTGTGGGCGCGGCGGTCGGGCTGTAACCCCCCCCTGCACCCCCCTCCCCGAGTTGCTGAGCACGGCCCGGCTTCGGGTGCGGGGCTCCGTGCGGGGCGTGGCGCGGGGCTCGCCGTGCCGGGCGGGGGGTGGCGGCAGGTGGGGGTGCCGGGCGGGGCGGGGCCGCCTCGGGCCGGGGAGGGCTCGGGGGAGGGGCGCGGCGGCCCCGGAGCGCCGGCGGCTGTCGAGGCGCGGCGAGCCGCAGCCATTGCCTTTTATGGTAATCGTGCGAGAGGGCGCAGGGACTTCCTTTGTCCCAAATCTGGCGGAGCCGAAATCTGGGAGGCGCCGCCGCACCCCCTCTAGCGGGCGCGGGCGAAGCGGTGCGGCGCCGGCAGGAAGGAAATGGGCGGGGAGGGCCTTCGTGCGTCGCCGCGCCGCCGTCCCCTTCTCCATCTCCAGCCTCGGGGCTGCCGCAGGGGGACGGCTGCCTTCGGGGGGGACGGGGCAGGGCGGGGTTCGGCTTCTGGCGTGTGACCGGCGGCTCTAGAGCCTCTGCTAACCATGTTCATGCCTTCTTCTTTTTCCTACAGCTCCTGGGCAACGTGCTGGTTGTTGTGCTGTCTCATCATTTTGGCAAAGAATTCACCATGGAAGACGCCAAAAACATAAAGAAAGGCCCGGCGCCATTCTATCCGCTAGAGGATGGAACCGCTGGAGAGCAACTGCATAAGGCTATGAAGAGATACGCCCTGGTTCCTGGAACAATTGCTTTTACAGATGCACATATCGAGGTGAACATCACGTACGCGGAATACTTCGAAATGTCCGTTCGGTTGGCAGAAGCTATGAAACGATATGGGCTGAATACAAATCACAGAATCGTCGTATGCAGTGAAAACTCTCTTCAATTCTTTATGCCGGTGTTGGGCGCGTTATTTATCGGAGTTGCAGTTGCGCCCGCGAACGACATTTATAATGAACGTGAATTGCTCAACAGTATGAACATTTCGCAGCCTACCGTAGTGTTTGTTTCCAAAAAGGGGTTGCAAAAAATTTTGAACGTGCAAAAAAAATTACCAATAATCCAGAAAATTATTATCATGGATTCTAAAACGGATTACCAGGGATTTCAGTCGATGTACACGTTCGTCACATCTCATCTACCTCCCGGTTTTAATGAATACGATTTTGTACCAGAGTCCTTTGATCGTGACAAAACAATTGCACTGATAATGAACTCCTCTGGATCTACTGGGTTACCTAAGGGTGTGGCCCTTCCGCATAGAACTGCCTGCGTCAGATTCTCGCATGCCAGAGATCCTATTTTTGGCAATCAAATCATTCCGGATACTGCGATTTTAAGTGTTGTTCCATTCCATCACGGTTTTGGAATGTTTACTACACTCGGATATTTGATATGTGGATTTCGAGTCGTCTTAATGTATAGATTTGAAGAAGAGCTGTTTTTACGATCCCTTCAGGATTACAAAATTCAAAGTGCGTTGCTAGTACCAACCCTATTTTCATTCTTCGCCAAAAGCACTCTGATTGACAAATACGATTTATCTAATTTACACGAAATTGCTTCTGGGGGCGCACCTCTTTCGAAAGAAGTCGGGGAGGCGGTTGCAAAACGCTTCCATCTTCCAGGGATACGACAAGGATATGGGCTCACTGAGACTACATCAGCTATTCTGATTACACCCGAGGGGGATGATAAACCGGGCGCGGTCGGTAAAGTTGTTCCATTTTTTGAAGCGAAGGTTGTGGATCTGGATACCGGGAAAACGCTGGGCGTTAATCAGAGAGGCGAATTATGTGTCAGAGGACCTATGATTATGTCCGGTTATGTAAACAATCCGGAAGCGACCAACGCCTTGATTGACAAGGATGGATGGCTACATTCTGGAGACATAGCTTACTGGGACGAAGACGAACACTTCTTCATAGTTGACCGCTTGAAGTCTTTAATTAAATACAAAGGATACCAGGTGGCCCCCGCTGAATTGGAGTCGATATTGTTACAACACCCCAACATCTTCGACGCGGGCGTGGCAGGTCTTCCCGACGATGACGCCGGTGAACTTCCCGCCGCCGTTGTTGTTTTGGAGCACGGAAAGACGATGACGGAAAAAGAGATCGTGGATTACGTCGCCAGTCAAGTAACAACCGCGAAAAAGTTGCGCGGAGGAGTTGTGTTTGTGGACGAAGTACCGAAAGGTCTTACCGGAAAACTCGACGCAAGAAAAATCAGAGAGATCCTCATAAAGGCCAAGAAGGGCGGAAAGTCCAAATTGTAAGGATCCTTTCTTTCTTTAAATAATTTCTACTAAGTGTAGATAATGCCTAGACCTGTTGGGAAAATAATTTCTACTAAGTGTAGATCTCCCTATCAGTGATAGAGAACAAATAATTTCTACTAAGTGTAGATCGAGTTTTCTTTTTGTTTTGTACAAGTAAGTTAACTTGTTTATTGCAGCTTATAATGGTTACAAATAAAGCAATAGCATCACAAATTTCACAAATAAAGCATTTTTTTCACTGCATTCTAGTTGTGGTTTGTCCAAACTCATCAATGTATCTTATCATGTCTGGATC

CAGp-BFP-Triplex-Leader-Array-Terminator (Fig. 4)

CAG promoter

BFP

Triplex

Lachnospiraceae bacterium leader sequence

CRISPR array (AAAT separator, Repeat)

SV40 terminator

GTGAGCCCCACGTTCTGCTTCACTCTCCCCATCTCCCCCCCCTCCCCACCCCCAATTTTGTATTTATTTATTTTTTAATTATTTTGTGCAGCGATGGGGGCGGGGGGGGGGGGGGCGCGCGCCAGGCGGGGCGGGGCGGGGCGAGGGGCGGGGCGGGGCGAGGCGGAGAGGTGCGGCGGCAGCCAATCAGAGCGGCGCGCTCCGAAAGTTTCCTTTTATGGCGAGGCGGCGGCGGCGGCGGCCCTATAAAAAGCGAAGCGCGCGGCGGGCGGGAGTCGCTGCGTTGCCTTCGCCCCGTGCCCCGCTCCGCGCCGCCTCGCGCCGCCCGCCCCGGCTCTGACTGACCGCGTTACTCCCACAGGTGAGCGGGCGGGACGGCCCTTCTCCTCCGGGCTGTAATTAGCGCTTGGTTTAATGACGGCTCGTTTCTTTTCTGTGGCTGCGTGAAAGCCTTAAAGGGCTCCGGGAGGGCCCTTTGTGCGGGGGGGAGCGGCTCGGGGGGTGCGTGCGTGTGTGTGTGCGTGGGGAGCGCCGCGTGCGGCCCGCGCTGCCCGGCGGCTGTGAGCGCTGCGGGCGCGGCGCGGGGCTTTGTGCGCTCCGCGTGTGCGCGAGGGGAGCGCGGCCGGGGGCGGTGCCCCGCGGTGCGGGGGGGCTGCGAGGGGAACAAAGGCTGCGTGCGGGGTGTGTGCGTGGGGGGGTGAGCAGGGGGTGTGGGCGCGGCGGTCGGGCTGTAACCCCCCCCTGCACCCCCCTCCCCGAGTTGCTGAGCACGGCCCGGCTTCGGGTGCGGGGCTCCGTGCGGGGCGTGGCGCGGGGCTCGCCGTGCCGGGCGGGGGGTGGCGGCAGGTGGGGGTGCCGGGCGGGGCGGGGCCGCCTCGGGCCGGGGAGGGCTCGGGGGAGGGGCGCGGCGGCCCCGGAGCGCCGGCGGCTGTCGAGGCGCGGCGAGCCGCAGCCATTGCCTTTTATGGTAATCGTGCGAGAGGGCGCAGGGACTTCCTTTGTCCCAAATCTGGCGGAGCCGAAATCTGGGAGGCGCCGCCGCACCCCCTCTAGCGGGCGCGGGCGAAGCGGTGCGGCGCCGGCAGGAAGGAAATGGGCGGGGAGGGCCTTCGTGCGTCGCCGCGCCGCCGTCCCCTTCTCCATCTCCAGCCTCGGGGCTGCCGCAGGGGGACGGCTGCCTTCGGGGGGGACGGGGCAGGGCGGGGTTCGGCTTCTGGCGTGTGACCGGCGGCTCTAGAGCCTCTGCTAACCATGTTCATGCCTTCTTCTTTTTCCTACAGCTCCTGGGCAACGTGCTGGTTGTTGTGCTGTCTCATCATTTTGGCAAAGAATTGAATTCGTCGCCACCATGGAGCTGATTAAGGAGAACATGCACATGAAGCTGTACATGGAGGGCACCGTGGACAACCATCACTTCAAGTGCACATCCGAGGGCGAAGGCAAGCCCTACGAGGGCACCCAGACCATGAGAATCAAGGTGGTCGAGGGCGGCCCTCTCCCCTTCGCCTTCGACATCCTGGCTACTAGCTTCCTCTACGGCAGCAAGACCTTCATCAACCACACCCAGGGCATCCCCGACTTCTTCAAGCAGTCCTTCCCTGAGGGCTTCACATGGGAGAGAGTCACCACATACGAAGACGGGGGCGTGCTGACCGCTACCCAGGACACCAGCCTCCAGGACGGCTGCCTCATCTACAACGTCAAGATCAGAGGGGTGAACTTCACATCCAACGGCCCTGTGATGCAGAAGAAAACACTCGGCTGGGAGGCCTTCACCGAGACGCTGTACCCCGCTGACGGCGGCCTGGAAGGCAGAAACGACATGGCCCTGAAGCTCGTGGGCGGGAGCCATCTGATCGCAAACATCAAGACCACATATAGATCCAAGAAACCCGCTAAGAACCTCAAGATGCCTGGCGTCTACTATGTGGACTACAGACTGGAAAGAATCAAGGAGGCCAACAACGAGACCTACGTCGAGCAGCACGAGGTGGCAGTGGCCAGATACTGCGACCTCCCTAGCAAACTGGGGCACAAGCTTAATTAAGAATTCGGATCACGCGTGATTCGTCAGTAGGGTTGTAAAGGTTTTTCTTTTCCTGAGAAAACAACCTTTTGTTTTCTCAGGTTTTGCTTTTTGGCCTTTCCCTAGCTTTAAAAAAAAAAAAGCAAAAGGTACCTGATTTAATTGCAAATCTTTGAAATAATGCAGACTTAAATTTATAAATTCATGGAATAAGGTGATTTTATTGTGAAAAAATACTCGTATTTTGTTGGAAAAACATCTTTTTGTTGTATAATATGATGATATACGGGATCCTTTCTTTCTTTAAATAATTTCTACTAAGTGTAGATAAAAGTGCCACTCCTTAGGGAAATAATTTCTACTAAGTGTAGATAGATGAGATGGTGACAGATAAAATAATTTCTACTAAGTGTAGATCTCCAGAGCCCGACTCGCCGAAATAATTTCTACTAAGTGTAGATTCGGCTTGCGGGGAGACTTCAAATAATTTCTACTAAGTGTAGATCACCCTTCGGCCCGCCACCCAAATAATTTCTACTAAGTGTAGATAGAAAGGGAATCCCAGGGCCAAATAATTTCTACTAAGTGTAGATCTCCCTATCAGTGATAGAGAAAATAATTTCTACTAAGTGTAGATGTTAACTTGTTTATTGCAGCTTATAATGGTTACAAATAAAGCAATAGCATCACAAATTTCACAAATAAAGCATTTTTTTCACTGCATTCTAGTTGTGGTTTGTCCAAACTCATCAATGTATCTTATCATGTCTGGATC
