## Supplemental Figures for "Enhanced Cas12a multi-gene regulation using a CRISPR array separator"

Figure S1

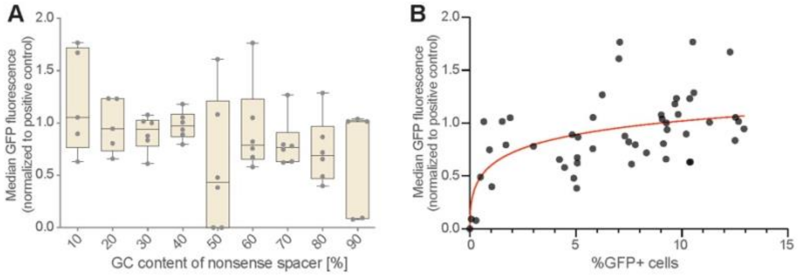


**Figure S1. CRISPR-activation of genomically integrated GFP**

(**A**) The negative correlation between GFP activation and nonsense-spacer GC content (**Fig. 1G**) is seen also when measuring median GFP fluorescence, though *percentage of cells that activate GFP* is a more sensitive measure of CRISPR activation effects than the *median GFP fluorescence* **(B)**, at least in this experimental system.

Figure S2

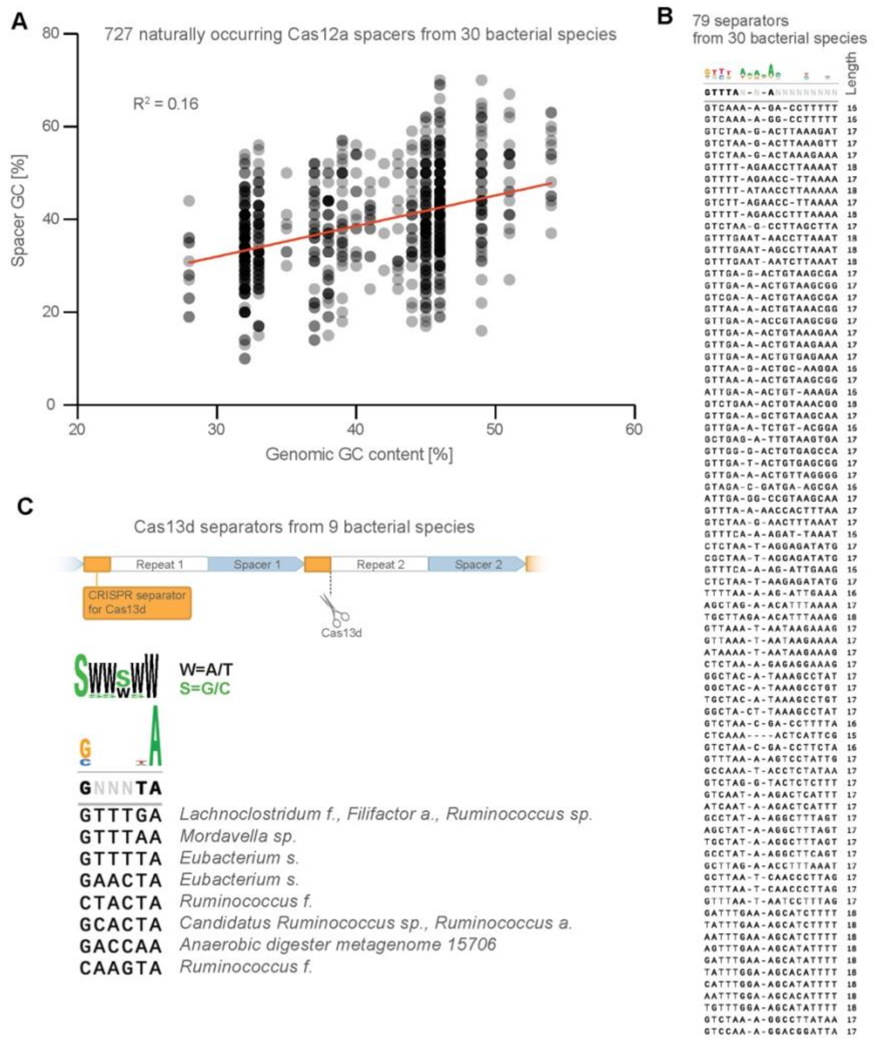


**Figure S2. Further analysis of naturally occurring crRNA sequences**

(**A**) The GC content of naturally occurring spacers is weakly correlated with overall genomic GC content of their bacterial host. (**B**) Full list of CRISPR-separator sequences from **Fig. 2C.** (**C**) CRISPR-separators from the type VI CRISPR Cas13d are also AT-rich, particularly the two bases at the very 3' end.

**Table S1. Spacer, repeat and separator sequences used and analyzed in this study**

**Table S2. Naturally occurring CRISPR-Cas sequences analyzed in this study**

**Table S3. Spacers used for endogenous gene activation (related to Fig. 4)**

**Table S4. Raw data used for figures in this study**

**Supplementary File 1. Sequences of expression constructs used in this study.**
